# The impact of structural variation on human gene expression

**DOI:** 10.1101/055962

**Authors:** Colby Chiang, Alexandra J. Scott, Joe R. Davis, Emily K. Tsang, Xin Li, Yungil Kim, Farhan N. Damani, Liron Ganel, GTEx Consortium, Stephen B. Montgomery, Alexis Battle, Donald F. Conrad, Ira M. Hall

## Abstract

Structural variants (SVs) are an important source of human genetic diversity but their contribution to traits, disease, and gene regulation remains unclear. The Genotype-Tissue Expression (GTEx) project presents an unprecedented opportunity to address this question due to the availability of deep whole genome sequencing (WGS) and multi-tissue RNA-seq data from 147 individuals. We used comprehensive methods to identify 24,157 high confidence SVs, and mapped cis expression quantitative trait loci (eQTLs) in 13 tissues via joint analysis of SVs, single nucleotide (SNV) and short insertion/deletion (indel) variants. We identified 24,801 eQTLs affecting the expression of 10,101 distinct genes. Based on haplotype structure and heritability partitioning, we estimate that SVs are the causal variant at 3.3-7.0% of eQTLs, which is nearly an order of magnitude higher than prior estimates from low coverage WGS and represents a 26- to 54-fold enrichment relative to their scarcity in the genome. Expression-altering SVs also have significantly larger effect sizes than SNVs and indels. We identified 787 putatively causal SVs predicted to directly alter gene expression, most of which (88.3%) are noncoding variants that show significant enrichment at enhancers and other regulatory elements. By evaluating linkage disequilibrium between SVs, SNVs and indels, we nominate 49 SVs as plausible causal variants at published genome-wide association study (GWAS) loci. Remarkably, 29.9% of the common SV-eQTLs are not well tagged by flanking SNVs, and we observe a notable abundance (relative to SNVs and indels) of rare, high impact SVs associated with aberrant expression of nearby genes. These results suggest that comprehensive WGS-based SV analyses will increase the power of both common and rare variant association studies.

---

Over the past decade, genome-wide association studies (GWAS) have linked thousands of common genetic variants to human traits and diseases. Fine-mapping causal variants at GWAS loci has proven difficult because the vast majority (~88%) reside in noncoding genomic regions, and in most cases the causal variant(s) and relevant gene(s) or functional element(s) are not known^1^. This has confounded the identification of therapeutic targets for precision medicine. To bridge the gap between molecular and clinical phenotypes, genome-wide eQTL scans have sought to identify genetic determinants of gene expression variation as markers of functional effect and a bridge connecting germline genetic variation to somatic cell biology^2–4^. These studies have successfully identified tens of thousands of eQTLs in a variety of human tissues.

A notable limitation of most extant eQTL studies is that, due to their reliance on SNP genotyping arrays, it has been difficult to identify the causal variants underlying eQTL associations and to judge the relative contribution of different variant classes to genetically regulated expression. Of particular interest is structural variation (SV), a broad class of variation that includes copy number variants (CNVs), balanced rearrangements and mobile element insertions (MEIs). Structural variation is recognized to be an important source of genetic diversity — 5,000 to 10,000 SVs are detectable in a typical human genome using short-read DNA sequencing technologies — but little is known about the mechanisms through which SVs affect gene expression and phenotypic variation. Although SVs are less abundant than SNVs, which represent ~4 million variant sites per genome^5^, SVs account for a greater number of nucleotide sequence differences due to their size, and may therefore exhibit outsized phenotypic effects^6,7^. Indeed, SVs have been identified as causal contributors to a number of rare and common diseases, and are generally presumed to act through their effects on gene expression^8^.

Despite many noteworthy examples linking SVs to gene expression and phenotypic variation in humans, more general and quantitative questions regarding the contribution of SVs relative to other variant classes remain a matter of debate. Two early studies used low-resolution microarray technologies to study the relationship between CNVs and gene expression, but their conclusions were limited to large CNVs that are now known to comprise a small fraction of SV^9,10^. A recent study from the 1000 Genomes Project represents the most comprehensive analysis to date, using RNA-seq expression profiles from lymphoblastoid cell lines (LCLs) of 462 individuals^2^ and SVs identified from low-coverage (median 7.4X) WGS data^11^. This analysis identified 9,591 eQTLs, of which 54 had an SV as the lead marker (denoted SV-eQTLs), implying that SVs are the causal variant at 0.56% of eQTLs. However, the study’s shallow sequencing depth limited SV detection power and genotyping accuracy, which are known to suffer in low-coverage data^12^. Furthermore, although gene expression is differentially regulated across tissues, prior SV-eQTL studies have focused solely on LCLs, and it is not known whether these observations extend to other cell types.

Here, we utilized multi-tissue RNA-seq expression data from the GTEx project to perform the first comprehensive human eQTL mapping study from deep WGS (median 49.9X) data that directly measures the contribution of SVs, SNVs, and indels.

We analyzed 147 human samples using SpeedSeq^13^ for alignment, data processing and per-sample SV breakpoint detection via LUMPY^12^, followed by cohort-level breakpoint merging, refinement, classification, and genotyping (Online Methods). We used complementary read-depth analysis with Genome STRiP to detect additional CNVs^14^. Together, these methods yielded a total of 45,968 structural variants, including a “high confidence” set of 24,157 SVs that met strict quality filters and are the basis for all subsequent analyses (**Table 1**). Several features indicate that these calls are high quality: we detected consistent numbers and proportions of SVs per sample (**Fig. 1a, b**), African American samples had an average of 27% more heterozygous LUMPY deletions than other samples (in accordance with previous observations^11^), the SV minor allele frequency (MAF) distribution mirrored that of SNVs and indels (Fig. 1c), and the SV size distribution is similar to prior WGS-based studies (**Fig. 1d, Supplementary Fig. 1**). This comprehensive variant call set is a powerful resource for functional analyses due to the high resolution (median breakpoint confidence interval: 39 bp) and diverse variant types including deletions (50.4%), duplications (14.7%), multi-allelic CNVs (mCNVs; 6.8%), mobile element insertions (MEIs; 8.5%), inversions (0.2%), and novel adjacencies of indeterminate type (hereafter denoted as “breakends”, or BNDs; 19.3%)^15^.

**Table 1.** Summary of variant types and discovery methods. SNVs and indels were detected using the Genome Analysis Toolkit (GATK) and SVs were detected by breakpoint evidence (BP), read-depth evidence (RD), or both. A subset of common variants were tested for *cis*-eQTLs, and were required to be on the autosomes or X chromosome and present in a minimum of 10 of the individuals with RNA-seq in at least one tissue. The SV-only eQTL mapping excluded SNVs and indels for greater sensitivity, while the joint eQTL mapping included all variant types. *Resolution refers to the positional certainty at each breakpoint, with read-depth variants having approximate breakpoint precision on the kilobase scale.

|  | Detection method | # of variants | Median resolution (bp) | Median size (bp) | # of common variants | SV-only eQTLs | Joint eQTLs |
| --- | --- | --- | --- | --- | --- | --- | --- |
| SNV | GATK | 21,764,904 | - | 1 | 6,982,921 | - | 17,024 (0.2%) |
| Indel | GATK | 3,030,964 | - | 3 | 834,255 | - | 2,245 (0.3%) |
| Deletion (DEL) | BP, RD | 11,692 | 34 | 966 | 3,373 | 546 (16.2%) | 27 (0.8%) |
|  | RD | 493 | kilobase* | 3,699 | 275 | 63 (22.9%) | 16 (5.8%) |
| Duplication (DUP) | BP, RD | 2,514 | 97 | 577 | 798 | 103 (12.9%) | 3 (0.4%) |
|  | RD | 1,054 | kilobase* | 4,993 | 833 | 148 (17.8%) | 70 (8.4%) |
| Multi-allelic CNV (mCNV) | RD | 1,635 | kilobase* | 3,676 | 992 | 241 (24.2%) | 100 (10.1%) |
| Inversion (INV) | BP | 58 | 10 | 573 | 19 | 2 (10.5%) | 0 (0.0%) |
| Mobile element insertion (MEI) | BP | 2,053 | 1 | 21 | 1,610 | 269 (16.7%) | 11 (1%) |
| Other SV (BND) | BP | 4,658 | 33 | - | 2,220 | 298 (13.4%) | 6 (0.7%) |
| All SVs | - | 24,157 | 178 | - | 10,120 | 1,670 (16.5%) | 233 (2.3%) |
| All variants | - | 24,820,025 | - | - | 7,827,296 | - | 19,269 (0.2%) |

We mapped *cis* eQTLs using 10,120 common SVs and whole transcriptome RNA-seq data from 13 tissues. We defined a *cis* window to include SVs within 1 Mb of each gene, and applied a permutation-based eQTL mapping approach using FastQTL, revealing 5,153 SV-eQTLs associated with expression differences at 2,102 distinct genes (hereafter referred to as “eGenes”) and 1,670 distinct SVs (Benjamini-Hochberg false discovery rate (FDR): 10%)^16^ (**Supplementary Table 1**).

SVs altered exons at 11.0% of eQTLs, providing a testable framework for their causal effects. Loss of function variants such as deletions or exon-disrupting MEIs are expected to decrease gene expression, exon duplications should increase gene expression, and neutral markers that tag a nearby causal variant through linkage disequilibrium (LD) should show bidirectional effects. Indeed, 513/568 (90.3%) of exon-altering eQTLs showed patterns of expression consistent with the SV class (**Fig. 2a**). This finding establishes strong evidence of a causal role for SVs at a subset of eGenes. In contrast, the remaining 4,585 eQTLs (89.0%) generally exhibited bidirectional expression effects (**Fig. 2a**). This may reflect a complex regulatory landscape of both enhancing and repressing DNA elements, or loci at which the SV is merely in LD with the true causal variant.

**Figure 1.**
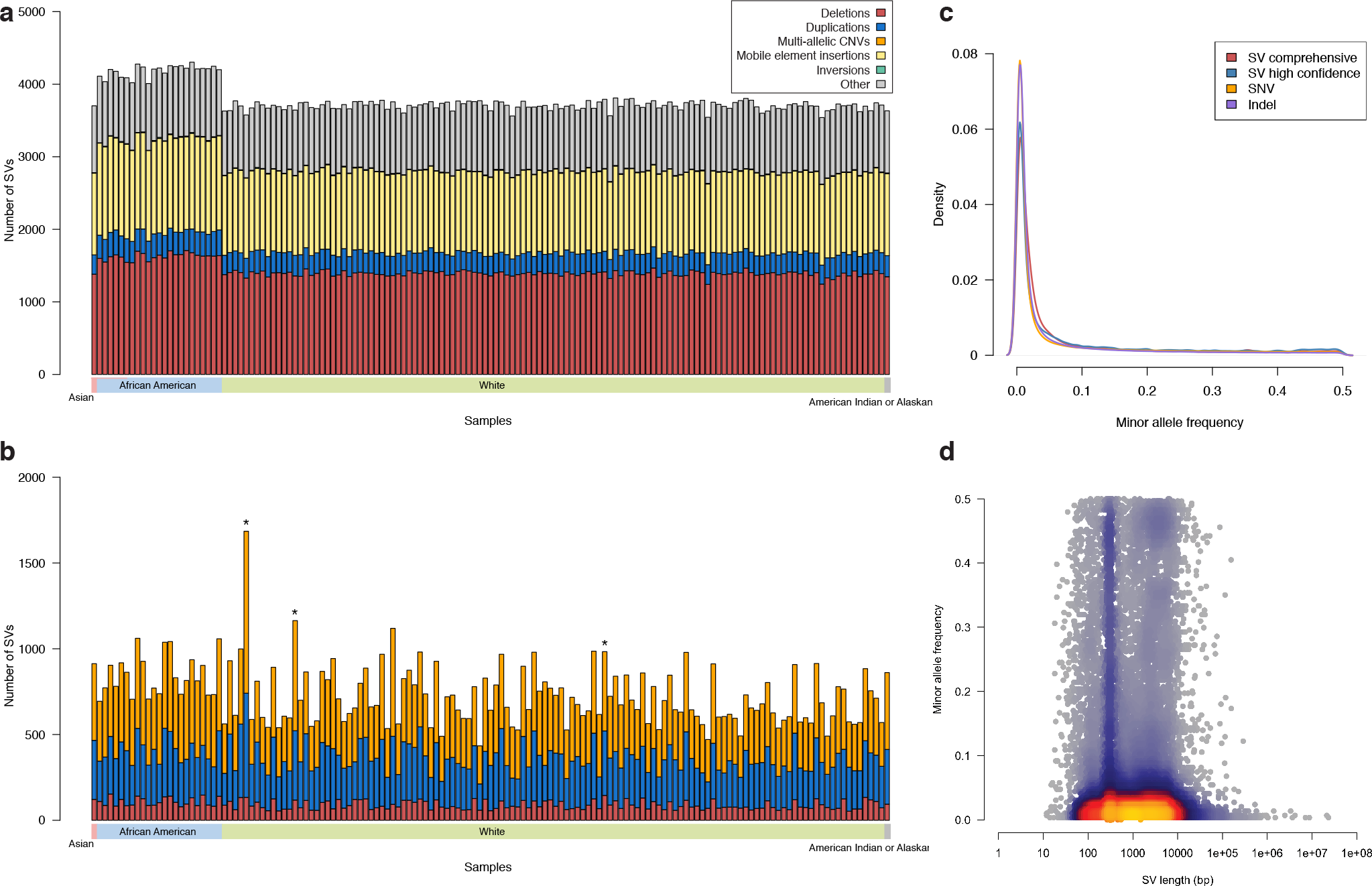
Structural variation call set. Number of SVs detected in sample by breakpoint evidence (with supporting read-depth evidence for deletions and duplications) (a), and by read-depth evidence alone (b) in 147 deep (30-50X) human whole genomes. Starred (*) samples exhibited abnormal read-depth profiles, and were excluded from rare variant analyses. (c) The minor allele frequency distribution of SVs mirrored that of high quality SNVs and indels detected by GATK. (d) Heat scatter plot showing the relationship between SV length and minor allele frequency, with a peak at ~300 bp due to Alu SINE insertions.

To assess the relative contribution of SV, we expanded our eQTL analysis to include 6,982,921 SNVs and 822,241 indels detected by the Genome Analysis Toolkit (GATK) ^17^. We performed joint eQTL mapping with the complete set of genetic variants, nominating a most likely causal variant for each eQTL identified. This produced 23,441 joint eQTLs affecting 9,547 distinct eGenes including 790 SV-eQTLs (3.4%), 20,208 SNV-eQTLs (86.2%), and 2,443 indel-eQTLs (10.4%). The observation that SVs are the lead marker at 3.4% of eQTLs provides an initial estimate of their contribution to gene expression variation. This is ~6-fold larger than a similar estimate from the 1000 Genomes Project, where merely 0.56% of eQTLs identified in LCLs had an SV as the lead marker^11^. These disparate results are not due to differences among tissues – we find that SVs are the lead marker at 2.8% of eQTLs identified in whole blood – and are unlikely to stem from trivial methodological differences given the similarity of eQTL mapping methods used in the two studies, and the fact that we recapitulated 31 of 47 (66.0%) previously identified LCL SV-eQTLs at eGenes also expressed in available tissues. We suspect that the key difference between these studies is the greater sensitivity and accuracy afforded by deep WGS data, which underscores the power and novelty of our study.

**Figure 2.**
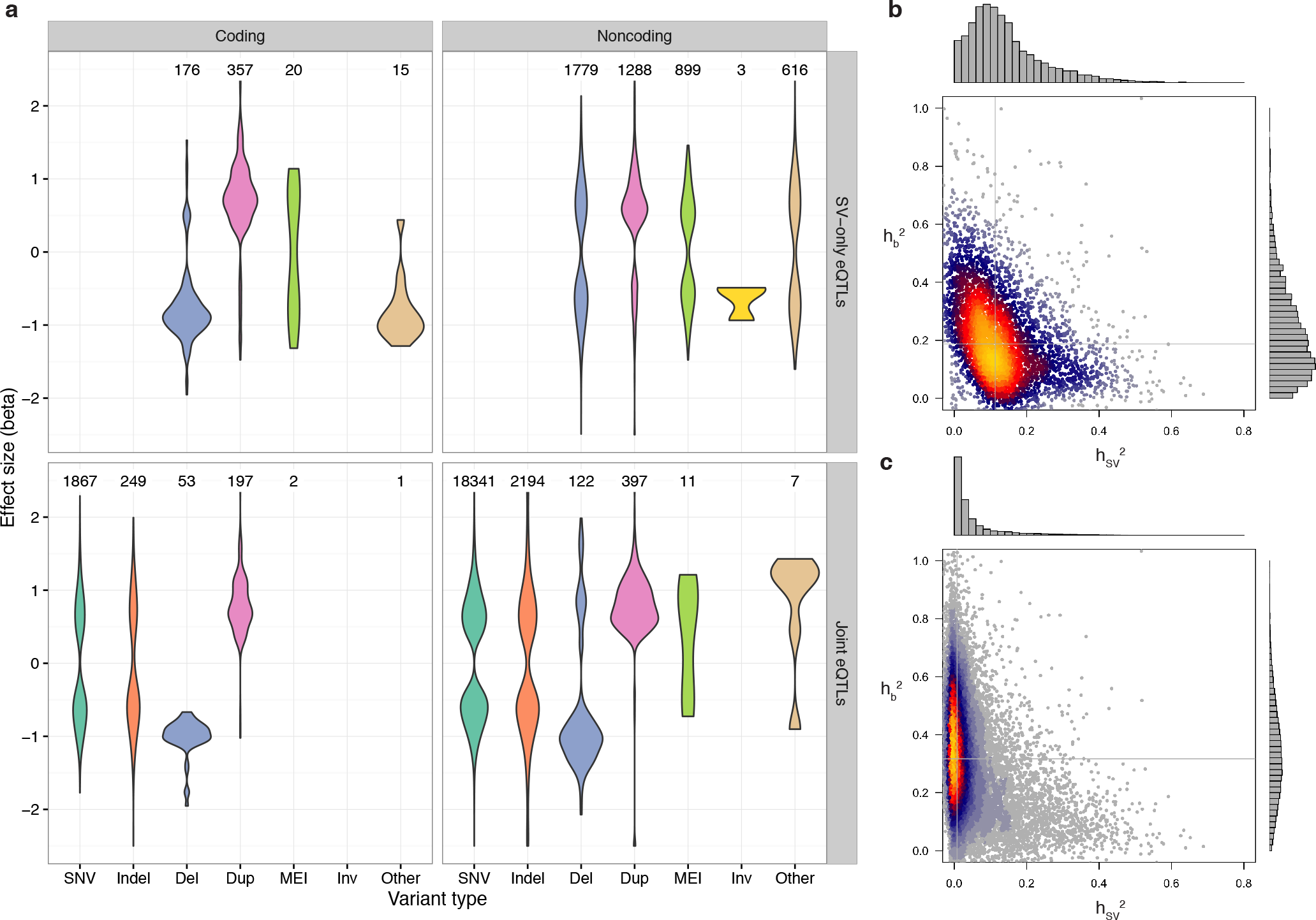
eQTL effect size distributions and heritability partitioning with linear mixed models. (**a**) Effect size distributions for coding and noncoding variants of each type, with the number of eQTLs of each type above each distribution. The top panels (SV-only eQTLs) show the 5,153 eQTLs that were discovered by the SV-only analysis, while the bottom two panels show the 23,441 eQTLs discovered by the joint analysis. Multi-allelic CNVs are included in the “Dup” category for this plot. (**b,c**) Heat scatter plots showing the heritability of each eQTL apportioned to the most significant SV in the *cis* window (x-axis) and the additive effect from the top 1,000 most significant SNVs and indels in the *cis* window (y-axis) for (**b**) SV-only and (**c**) joint eQTL mapping analyses. Gray lines mark the median of values for each axis.

We next applied fine-mapping approaches to infer the probability that each locus contained a causal SV in the eGene’s *cis* window. At each of the 23,441 joint eQTLs, we identified the 100 SNVs and indels in the 1 Mb *cis* window that were most significantly associated with the eGene’s expression by their FastQTL nominal p-value, as well as the single most significant SV. We then used the CAVIAR software package to apportion a causal likelihood and a relative ranking to each of these 101 markers based on the magnitude and direction of association as well as the pairwise LD structure across the region^18^. This approach aims to disentangle each variant’s causal contribution from its association due to LD with nearby causal markers. At 3.3% of eQTLs overall (2.2-4.3% among tissues), the SV was identified among the 101 candidates as the highest probability causal variant underlying the eQTL association.

As an orthogonal estimate of contribution of SV, we applied a linear mixed model to partition the heritability of each eGene’s expression into a fixed effect from the SV, and a random effect representing the cumulative heritability of the 1,000 most significant SNPs and indels in the *cis* region (**Fig 2b,c, Supplementary Fig. 2**). This method mirrors that of several prior studies that have examined relative contributions of distinct variant classes on a quantitative trait^19–21^. Heritability partitioning revealed that SVs account for 8.5% of total gene expression heritability when summing their effects across all eQTLs, although we note that this includes numerous loci where the SV has a very small effect. More importantly, at the 22,289 eQTLs that showed appreciable overall genetic heritability (>0.05), the SV contributed more heritability than the additive effect of the other 1,000 variants in 7.0% of cases, suggesting that the SV was the causal variant.

Taken together, the three independent analyses presented above – FastQTL-based eQTL mapping, CAVIAR-based fine mapping and GCTA-based heritability estimation – indicate that SVs are the causal variant at 3.4%, 3.3% and 7.0% of eQTLs, respectively. These are likely to be underestimates because the genotyping error rate for SVs is higher than for SNVs and indels, giving the latter a relative advantage to “win” causal variant prediction tests in regions of strong LD. Although the absolute contribution of SVs to heritable expression variation is small compared to SNVs and indels, on a per-variant basis, an SV is 26-to 54-times more likely to modulate expression than an SNV or an indel. Moreover, SVs showed a 1.2-fold larger median effect size on gene expression than SNVs and indels (p-value: < 1 × 10^−15^, Mann-Whitney U test), and deletions showed a 1.4-fold larger median effect size, with direction of effect predominantly correlating with SV type (**Fig. 2a**). This result is unlikely to stem from differences in statistical power given the observed allele frequency distributions of each variant class, and the fact that SVs have consistently greater effect sizes across matched allele frequency bins (**Fig 1c, Supplementary Fig. 3**). Together, these results demonstrate that SVs play an important and outsized role in defining the landscape of genetically regulated gene expression.

We next sought to examine the genomic context of SV-eQTLs for clues into their molecular mechanisms. We hypothesized that causal SVs would be enriched in functional elements such as gene bodies, enhancers and repressors. To maximize the number of causal variants in this analysis, we first created an aggregate eQTL set containing the union of all eQTLs identified by either the SV-only or joint eQTL mapping (24,594 eQTLs affecting 10,094 distinct eGenes). We then derived a composite “causality score” that incorporates the aforementioned CAVIAR and GCTA estimates of SV causality at each eQTL by multiplying the CAVIAR posterior causal probability with the SV’s *cis* heritability fraction 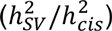 (**Supplementary Fig. 4**). At each eGene we selected the SV within 1 Mb that had the strongest association to the eGene’s expression, and allocated these 4,485 distinct SVs into seven bins according to their composite score quantile, with the least causal bin comprising the bottom half of composite scores, and the most causal bin comprising the 95th percentile. Different SV classes were represented in roughly consistent proportions across the lower causality bins, but the most causal bin of SVs had higher concentrations of multi-allelic CNVs and duplications (**Fig. 3a**).

**Figure 3.**
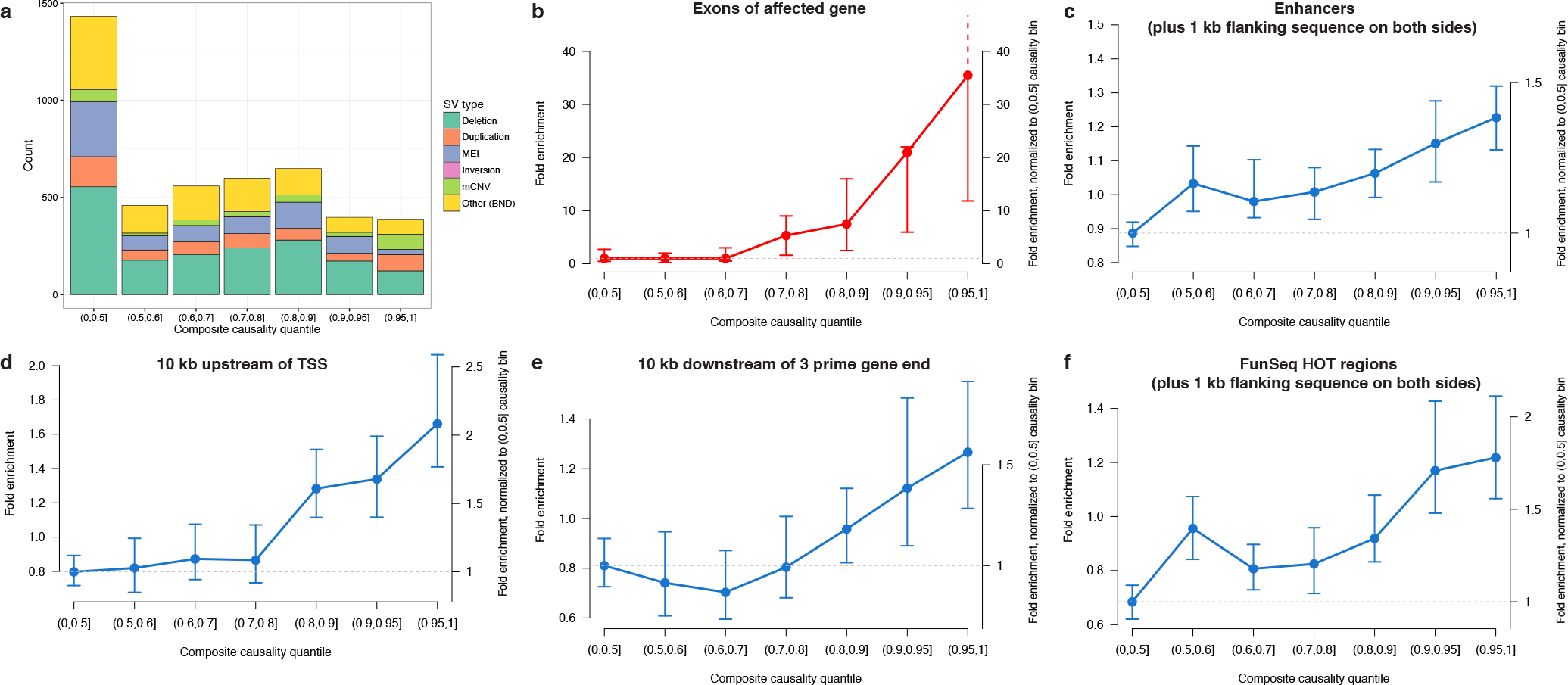
Feature enrichment of SV-eQTLs. Fold enrichment and 95% confidence intervals (based on 100 random shuffled sets of the positions of SVs in each bin) for the overlap between the most significant SV and various annotated genomic features at the union of eQTLs discovered by SV-only or joint eQTL mapping. (**a**) Composition of each causality score bin by SV type. (**b**) Enrichment for an SV in each bin of causality to touch exons of the affected eGene (**c-f**) For the remaining plots in blue, SVs that overlapped with an exon of their affected eGene were excluded, yet the remaining SVs still showed significant enrichment in enhancers (**c**), in the 10 kb regions upstream (**d**) and downstream of transcriptions start sites (TSS) (**e**), and regions predicted to be highly occupied by transcription factors (FunSeq HOT regions) (**f**).

We examined the overlap between SVs and annotated genomic features to assess enrichment in various functional elements. SVs in the 95th percentile of causality showed a 35.5-fold enrichment for altering eGene exons, amounting to 18.3% (71/389) of the most causal SVs, compared to 0.3% of SVs in the least causal bin representing the lower half of causality scores (**Fig. 3b**). Recapitulating the trend from SV-alone eQTL mapping (**Fig. 2a**), the expression effect direction was highly correlated with SV type (74/84 showing the expected direction), strongly suggesting that this set of exon-altering SVs are the causal variant at their respective eQTLs. Importantly, this analysis also demonstrates that our causality score effectively distinguishes neutral from causal SVs: no significant enrichment of exon-altering SVs is observed in bins beneath the 80th percentile of scores, and enrichment rises precipitously from the 80th to the 95th percentile.

However, the vast majority of SVs – including 81.7% of those in the 95th percentile of predicted causality – do not alter eGene dosage or structure, and thus are likely to act through regulatory mechanisms. We analyzed these 4,363 non-coding SVs for enrichment in other functional elements of the genome with potential regulatory consequences. We found that several functional elements were stratified by causality score and significantly enriched in the most causal bins, including the regions within 1 kb of enhancers, the regions 10 kb upstream or downstream of gene transcripts, and regions predicted by FunSeq to be highly occupied by transcription factors^22–24^ (**Fig. 3c–f, Supplementary Fig. 5**). In all cases, regulatory element enrichment was most pronounced in the top causality score bin – providing further evidence of the effectiveness of our scoring method – yet more moderate enrichments were also observed in lower bins. This demonstrates that SVs with strong causality predictions are more likely to affect known regulatory elements, strongly suggesting that many are *bona fide* causal variants that alter gene expression through regulatory effects. Given the extreme paucity of causal regulatory variants discovered in the human genome to date, the set of 695 promising candidates from the top 10% of causality scores will prove invaluable for future studies.

A major bottleneck for interpreting GWAS results is identifying causal variants and determining the genes and molecular mechanisms contributing to disease. Bridging this gap is a driving motivation for eQTL studies. Approximately 88% of GWAS loci are in noncoding regions of the genome^1^, suggesting that they act through effects on gene regulation. Indeed, ~6% of GWAS loci were found to be in strong LD (R^2^≥0.8) with eQTLs in a recent GTEx study using SNP arrays^3^ (7.7% with the WGS data described here), and the true number of disease-associated eQTLs is believed to be higher^25^. Thus, the SV-eQTLs identified here provide an unparalleled resource to assess the contribution of SV to common disease.

To investigate the contribution of SVs to trait-associated loci, we first identified 4,874 SNPs from the GWAS catalog that were non-redundant on a per-locus and per-disease basis, were genotyped in the GTEx samples, and that had convincing evidence for disease association (p<5×10^−8^) ^26^. Of these, 842 were in LD (R^2^≥0.5) with a lead marker from our joint SV/SNP/indel eQTL analysis, suggesting that the GWAS hit and the eQTL are produced by the same underlying causal variant. An SV was the candidate causal variant at 3.2%, 3.3% and 14.3% of the 842 GWAS-associated eQTLs, depending on whether causality is judged based on FastQTL, CAVIAR or GCTA (as in the prior causal SV analysis).

Combined with the eQTL fine mapping results presented above, this suggests that SVs underlie a similar fraction GWAS-associated eQTLs as they do eQTL results on the whole, indicating that our results are directly relevant to common disease biology.

We next screened for SVs that were likely to explain prior GWAS results. We identified 49 SVs in LD (R^2^≥0.5) with GWAS loci that were predicted to be the causal variant for an eQTL (**Supplementary Table 2**). Here, we define causal SVs as those in the top 10% of composite causality scores, a set that shows significant enrichment with functional annotations (**Fig. 3**). Ultimately, experimental validation will be required to definitively establish the causal relationship between any given variant and GWAS result. However, there are a number of promising candidates among these 49 loci. In one case, a 294 bp deletion is associated with decreased expression of the DAB2IP gene in thyroid tissue – apparently by disrupting an intronic enhancer – and is linked to a risk allele for abdominal aortic aneurysm^27^ (R^2^=0.57; **Fig 4a**). In another case, a 1,468 bp deletion in intron 10 of the PADI4 gene is linked (R^2^=0.70) to a risk allele for rheumatoid arthritis^28^ (**Fig. 4b**). Multiple studies have reported significant association between haplotypes of the PADI4 gene and rheumatoid arthritis^29,30^, and high levels of PADI4 mRNA have been detected in pathological synovial tissues, yet none have implicated this deletion, which flanks an annotated enhancer and is predicted to be the causal variant for increased PADI4 expression in lung. A third example relates an SVA retroelement insertion to a GWAS risk allele for melanoma and esophageal cancer (R^2^=0.85) ^31,32^ (**Fig. 4c**). The variant is a polymorphic SVA insertion that defines the boundaries of alternative splice isoforms of CASP8. The insertion allele is associated with reduced expression of CASP8, which encodes a protease with a critical role in the regulation of proliferation and apoptotic cell death (**Fig. 4d**). Finally, we recapitulate several SVs previously recognized as clinically associated markers, such as a ~32 kb deletion conferring risk for psoriasis^33^ (**Fig. 4d**) and a ~37 kb deletion linked to circulating liver enzyme levels (gamma-glutamyl transferase) ^34^ (**Supplementary Fig. 6**).

**Figure 4.**
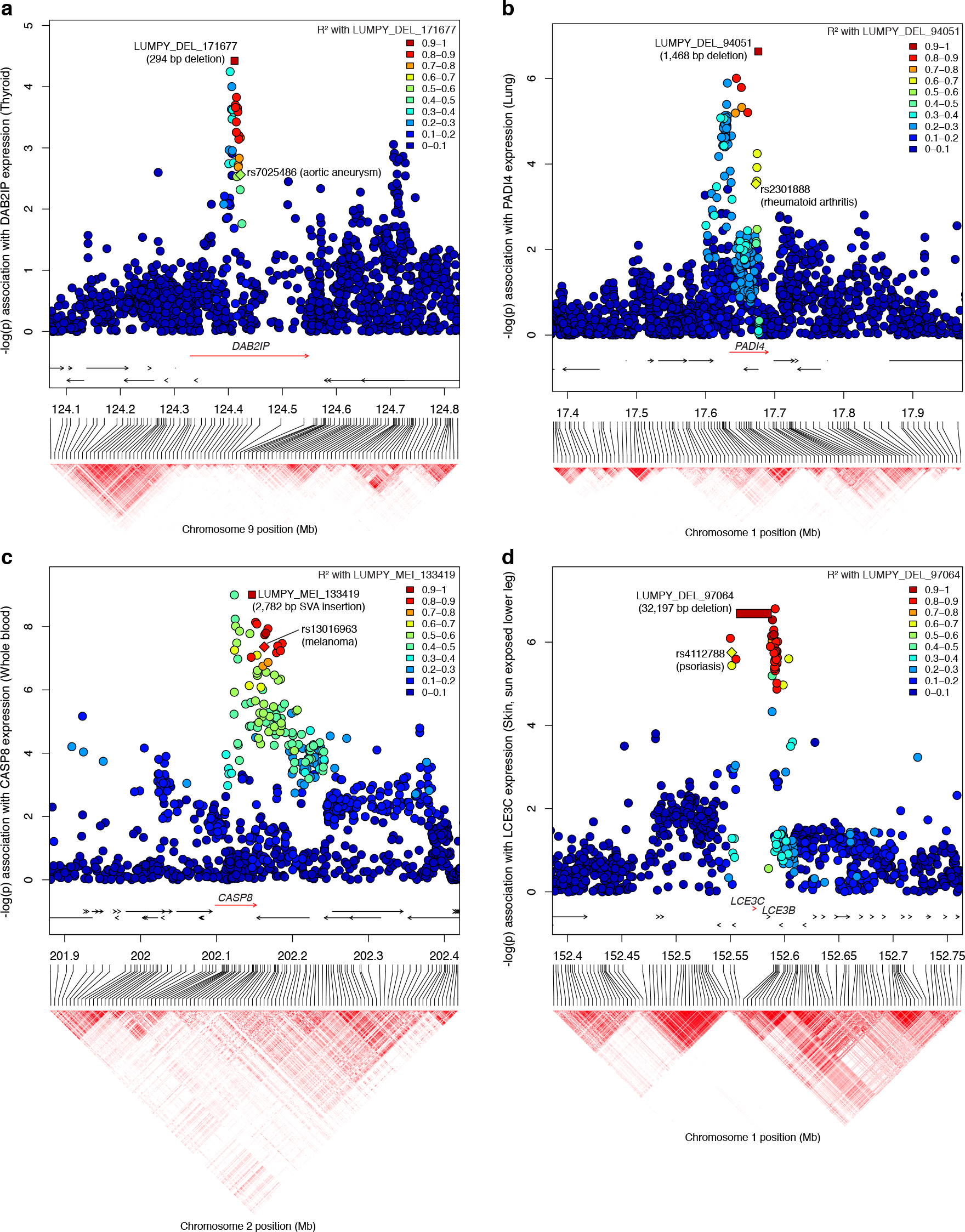
Candidate SV-eQTLs at GWAS loci. Genomic position and haplotype block are shown on the x-axis, and each variant’s association with the indicated eGene is shown on the y-axis. The rectangular points represent the predicted causal SV, with the colors representing its linkage (R^2^) to each marker in the window. The labeled diamond shows the reported risk allele for the specified GWAS phenotype (**a**) A 294 bp deletion that intersects an enhancer in intron 1 of DAB2IP was linked to a risk allele for abdominal aortic aneurysm (rs7025486), and is also predicted to be a causal eQTL for DAB2IP. (**b**) A 1,468 bp deletion associated with increased expression of PADI4 is linked to a known risk allele for rheumatoid arthritis (rs2301888). (**c**) A polymorphic mobile element insertion defining exon boundaries of CASP8 reduces the gene’s expression and is linked with a risk allele for melanoma (rs13016963). (**d**) A large 32,197 bp deletion of the LCE3C and LCE3B genes that was previously identified as a risk factor for psoriasis was recapitulated by our study.

However, GWAS based on SNP array genotypes can only inform eQTLs that are well tagged by a SNP on the genotyping array. Notably 30.3% of common SVs, as well as 29.9% of those in the top 10% of causality scores, lacked a SNP marker on the Illumina Omni 2.5 array in linkage disequilibrium (R^2^≥0.5), indicating that a substantial portion of SV-eQTLs are untested in typical disease association studies. In comparison, only 10.1% of SNP and 17.2% of indel joint eQTLs are not well tagged.

We next sought to assess the role of rare SVs on gene expression variation. In contrast to common variant eQTLs, which are caused by ancient mutations that have been subjected to natural selection, most rare variants arose recently and are more likely to have larger effect sizes and deleterious consequences. Rare variants are difficult to study via traditional eQTL approaches because any given variant is observed too infrequently within a set of samples to establish a statistical relationship with gene expression^35^. However, the effect of rare variants on gene expression can be assessed indirectly via bulk outlier enrichment analyses^36^. We thus identified 5,047 gene expression outliers (median: 30 per person; range: 10-298) in which an individual exhibited aberrant transcript dosage compared to the data set as a whole (Online Methods). Next, we identified 5,660,254 rare variants (4,691 SVs, 4,830,727 SNVs, and 824,836 indels) that were positively genotyped in at most two individuals. To reduce the effects of population stratification, we limited this analysis to the 117 Caucasian individuals with RNA-seq data in at least 5 tissues.

Rare variants were significantly enriched by 1.2-fold (95% CI: 1.2-1.3) within the gene body and the 5 kb flanking sequence of expression outliers (**Fig. 5a, Supplementary Table 3**). This enrichment is most pronounced for SVs (16.1-fold, 95% CI: 11.5-25.4), in which 355/5,047 (7.0%) of gene expression outliers harbored a rare SV compared to the null expectation of 22/5,047 (0.4%) in 1,000 random permutations of the sample expression values. Notably, expression-altering SVs were significantly larger than rare SVs on the whole (p-value: < 1 × 10^−15^, Mann-Whitney U test), and duplications were disproportionately represented (**Fig. 5b,c**). In several cases a single large SV caused multiple gene expression outliers: a 21.3 Mb duplication event was associated with 161 outliers within the region, and two large duplications (4.1 Mb and 2.5 Mb) were associated with 11 and 30 outliers, respectively. However, the enrichment of rare SVs around outlier genes is not driven by a handful of large events, since the majority of outlier-associated SVs (56/99) were only associated with a single gene (**Supplementary Fig. 7**). Moreover, on a per-variant basis, 99/4,691 (2.1%) of the rare SVs had an expression outlier within 5 kb compared to 10/4,691 (0.2%) in the permutation set, representing a 9.9-fold enrichment (95% CI: 5.8-19.8) (**Fig. 5b**). These findings demonstrate that rare SVs are a common cause of aberrantly expressed genes, contributing a median of approximately 1 gene expression outlier per person. We expect this to be a large underestimate given the strict definition of expression outliers used in this study – rare variants are likely to contribute to more modest changes in expression as well.

**Figure 5.**
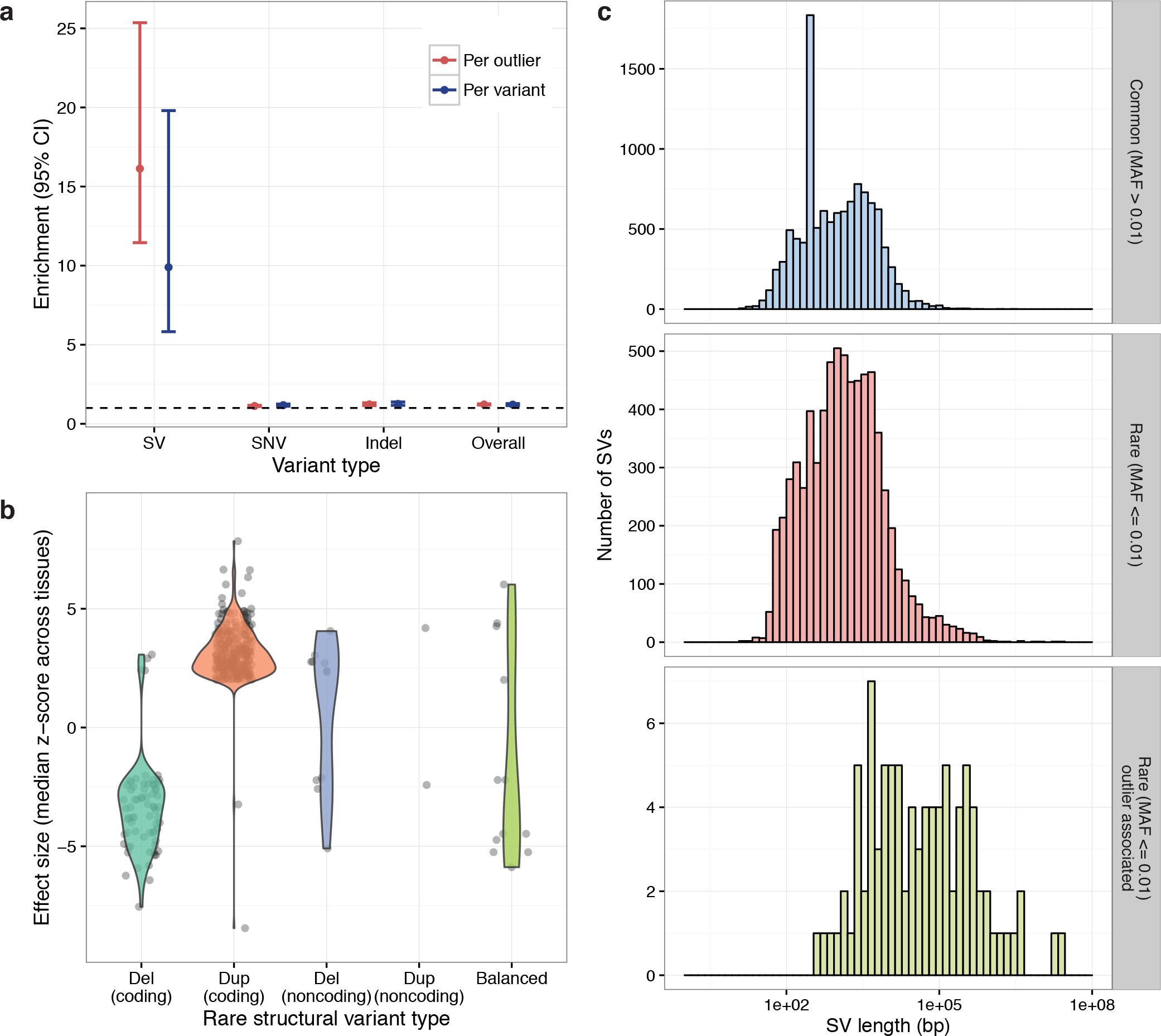
Gene expression outliers are associated with rare SVs. (**a**) Fold enrichment of rare variants within 5 kb of expression outliers (red) and fold enrichment of outliers within 5 kb of rare variants (blue) between the observed set of 5,047 outliers and 1,000 random permutations of their sample names. (**b**) Effect size distributions for each SV type within 5 kb of an outlier in the same individual, with “coding” SVs defined as those that overlap with exons of the outlier gene and “noncoding” defined by the remainder. (**c**) Size distribution histograms by minor allele frequency (MAF) for common (MAF>0.01), rare (MAF≤0.01), and rare SVs within 5 kb of an expression outlier in the same individual, excluding balanced rearrangements. A peak at ~300 bp in the topmost plot results from common Alu SINE insertions.

Our data show that rare SVs alter gene expression through diverse mechanisms. Of the 99 rare SVs predicted to causally alter gene expression (permutation-based FDR: 0.2%), 79 (79.8%) are CNVs that change dosage of the aberrantly expressed gene (**Supplementary Table 4**). Most gene expression changes occur in the expected direction relative to the dosage alteration (**Fig. 5c**), but we observed 4 deletions and 2 duplications with expression effects in the opposite direction; all involve partial gene alterations, which suggests complex regulatory effects rather than simple dosage compensation. The next most common class (11, 11.1%) are non-coding CNVs that appear to act through regulatory effects and – as in the case of the SV-eQTLs (**Fig 2a**) – show bidirectional effects on transcription. Remarkably, we identified a number of atypical SVs with strong yet unpredicted effects on gene expression. These include a 3.6 Mb inversion that alters the expression of 3 genes found at or near the breakpoints (one with increased and two with decreased expression), a 391 bp intronic inversion that causes increased expression, a complex 3-breakpoint balanced rearrangement that causes decreased gene expression, and 9 complex CNVs involving a combination of multiple copy number variable segments and/or adjacent balanced rearrangements, including one highly complex 6-breakpoint event that resembles chromothripsis (**Supplementary Table 5**). These results are consistent with prior studies describing the prevalence of complex SVs in “normal” human genomes, and reveal for the first time the diversity of gene expression effects caused by rare complex SVs.

We compared the relative contribution of rare SVs, SNVs and indels to expression outliers. Although the overall enrichment of SNVs and indels at gene expression outliers is mild due to the high background prevalence of rare variants in these classes, enrichment increases dramatically when analyses are restricted to high impact mutations (as judged by CADD^37^; **Supplementary Fig. 8**). Overall, we observed a net excess of 443 outliers within 5 kb of a rare variant in the same individual compared to the expected number from permutation tests, or 8.8% (443/5,047) of total outliers. Moreover, by partitioning excess outliers among SVs, SNVs and indels, we estimate that 69.9% of gene expression outliers with a genetic basis are likely explained by structural variation, whereas merely 15.9% and 14.4% are due to SNVs and indels, respectively (Online Methods, **Supplementary Fig. 9**). We note that this approximation assumes similar proportions of causal variants for each variant type, so may under-estimate the contribution of SNVs and indels. It also only captures the effects of rare variants within 5 kb of the outlier gene and depends on our definition of expression outliers. While the strength of the SV effect is due in part to 8 very large CNVs (> 1 Mb), GTEx individuals should be representative of the general population in terms of the prevalence of large CNVs, and the relative contribution of SVs remains noteworthy even when individuals with megabase-scale CNVs are excluded from the analysis (SV: 37.9%, SNV: 38.0%, indel: 24.1%). These results indicate that rare structural variants are a major cause of gene expression outliers in the human population, and suggest that thorough ascertainment of SV will significantly increase the power of rare variant association studies, and the efficacy of WGS-based disease diagnosis.

As human genetics moves deeper into the era of whole genome sequencing, it has become possible to include all forms of genetic variation in cohort studies and clinical practices. Our results demonstrate that comprehensive analysis of structural variation will be a critical aspect of these efforts.

## Online methods

### SV call set generation

We acquired 148 deep whole genome BAM files from the GTEx V6 data release (dbGaP accession phs000424.v6.p1). We excluded one sample (GTEX-WHWD-0002) due to an abnormal insert size distribution, which confounds SV detection. We realigned the remaining 147 whole genomes to GRCh build 37 plus a contig for Epstein-Barr virus using SpeedSeq v0.0.3 (BWA-MEM v0.7.10-r789) according to published practices^13,38^. We ran LUMPY v0.2.9 on each sample using the default parameters in the LUMPY Express script, using the published list of excluded genomic regions from SpeedSeq as well as the -P option to output probability curves for each breakpoint^12^. We merged the 147 VCF files using the l_sort.py and l_merge.py scripts included in LUMPY with the “--product” option and 20 bp of slop, simultaneously combining variants with overlapping breakpoint intervals while refining their spatial precision based on the probability curves to create a cohort-level VCF. We pruned remaining variants with nearly overlapping breakpoint intervals by selecting the single variant with the highest allele frequency among the overlapping set. Next, we genotyped each sample with SVTyper (https://github.com/hall-lab/svtyper), which performs breakpoint sequencing of paired-end and split-read discordants^13^. We define the term “allele balance” as the ratio of non-reference to total reads at each breakpoint. Allele balance serves a proxy for genotype that is tolerant to inefficiencies in aligning the alternate allele for SVs, and is used for most analyses in this paper. We then used CNVnator to annotate the copy number of each spanning variant (putative deletions, duplications, and inversions).

We applied several filters to the LUMPY call set to flag low quality SVs. Since 68 samples were sequenced on the Illumina HiSeq 2000 platform and 79 on the Illumina HiSeq X Ten platform, we flagged variants whose linear correlation (R^2^) between genotype and sequencing platform exceeded 0.1. We further flagged deletions lacking split-read support that were smaller than 418 bp, which was measured to be the empirical minimum deletion size at which all insert size libraries were able to discriminate between concordant and discordant reads with 95% certainty. We determined that three samples (GTEX-NPJ8-0004, GTEX-T2IS-0002, GTEX-0IZI-1026) had abnormal read-depth profiles and we therefore flagged SVs private to any of those samples, effectively excluding them from rare variant analyses. Finally, we flagged variants with a mean sample quality (MSQ, a measure of genotype quality among positively genotyped samples that is independent of allele frequency) of less than 20 as low quality.

Next, we reclassified variant types, requiring that deletions and duplications showed correlation between read-depth and the allele balance at the breakpoint. For common SVs (at least 10 samples with non-reference genotypes) we fit a linear regression and required a slope of at least 1.0 in the appropriate direction (positive for duplications, negative for deletions) and R^2^ ≥ 0.2. For the remaining low frequency SVs we required that > 50% of positively genotyped samples must be > 2 MAD (median absolute deviation) (in the correct direction for deletion/duplication) and > 0.5 absolute copies from the median of reference genotyped samples. For low frequency SVs on the sex chromosomes, we limited the above criteria to the gender with more individuals to avoid gender confounders. We identified mobile elements insertions (MEIs) in the reference genome as SVs with breakpoint orientations indicative of deletions that had > 0.9 reciprocal overlap with an annotated SINE, LINE, or SVA element with sequence divergence of less than 200 milliDiv, based on RepeatMasker annotations. Due to limitations of our pipeline, we were only able to detect MEIs inserted into the reference genome based on their absence in other genotyped samples.

We ran Genome STRiP 2.0 according to the best practices workflow for deeply sequenced genomes, using a window size of 1000 bp, window overlap of 500 bp, reference gap length of 1000 bp, boundary precision of 100 bp, and minimum refined length of 500 bp. We flagged CNVs for platform bias and the three samples with abnormal coverage profiles as described above. For rare SVs detected by Genome STRiP (private or doubletons in our call set) we merged fragmented variants with identical genotypes within 10 Mb of each other whose combined footprint encompassed at least 10% of their span.

We then unified the LUMPY and Genome STRiP call sets while collapsing redundancies. Because LUMPY variants are substantially more precise and have well-defined confidence intervals, we retained LUMPY calls when an SV was detected by both algorithms with a reciprocal overlap of > 0.5 and a matching variant type (mCNVs were allowed to merge with either LUMPY duplications or LUMPY deletions). To ensure that SVs would be merged even when the Genome STRiP call was fragmented, which occurs fairly often with GTEx WGS data, we also merged calls where > 0.9 of a Genome STRiP CNV was contained in a LUMPY SV of the same type (or mCNV) and their correlation between LUMPY allele balance and copy number had R^2^ > 0.25. This last step ensures that the merged variants have a high degree of co-occurrence among samples, and are not simply independent variants that inhabit the same genomic interval.

We measured allele frequency for LUMPY SVs as the ratio of non-reference to total alleles in the population. For Genome STRiP variants we defined allele frequency as the fraction of samples that deviate from the mode copy number value in the population.

We defined a high confidence call set from the variants that had not been flagged by the aforementioned filters. In general, high confidence SVs had to be supported by multiple independent evidence types. Since LUMPY deletions and duplications were identified by paired-end and/or split-read evidence and had also met requisite read-depth support from reclassification they were automatically considered high confidence. Similarly, MEIs were considered high confidence based on support from reference genome repeat annotations. LUMPY inversions and BNDs (unclassified breakends) were required to have a minimum variant quality score of 100. Inversions were further required to show evidence from both sides of the event, and at least 10% of supporting reads derived from each of split-read and paired-end evidence types. LUMPY BND variants were required to have at least 25% of supporting reads derived from each of split-read and paired-end evidence types. Genome STRiP variants that were merged as described above and those with GSCNQUAL score >=10 were considered high confidence. This set of 24,157 variants served as the basis for all analyses in this paper.

### Common eQTL mapping

We mapped *cis*-eQTLs to scan for significant associations between common variant genotypes and gene expression in all tissues for which there were ≥ 70 individuals with both WGS data and RNA-seq data. These include the following 12 tissues: whole blood, skeletal muscle, lung, tibial artery, aortic artery, adipose (subcutaneous), thyroid, esophagus mucosa, esophagus muscularis, skin (sun-exposed), tibial nerve, muscle (skeletal), as well as transformed fibroblasts. For convenience, we refer to the transformed fibroblasts as a tissue type throughout this study. RNA-seq data from each tissue was aligned with Tophat v1.4 using GENCODE v19 gene annotations by taking the union of exons for gene level quantification, and RPKM values were calculated with RNA-SeQC^39–41^. Reads were required to align exclusively within exons or span them (without aligning to intronic regions), align in proper pairs, contain a maximum of six non-reference bases, and map uniquely to the gene. Samples were quantile normalized within each tissue followed by inverse quantile normalization of each gene to control outliers.

We selected common genetic markers for which there were at least 10 samples with minor allele genotypes in our cohort. For Genome STRiP CNVs, we defined the reference genotype as the mode copy number observed in our data set. We performed two independent *cis*-eQTL mapping runs. The first, an “SV-only” eQTL analysis, used only common SV markers as genotype input for improved sensitivity under a reduced multiple-testing burden. The second, a “joint” eQTL analysis, included the 10,120 common SVs as well as 6,982,921 SNVs and 834,255 indels detected by the Genome Analysis Toolkit HaplotypeCaller v3.1-144-g00f68a3^17^, allowing a fair comparison of the relative contribution of different variant types.

We mapped cis-eQTLs with FastQTL v2.184 using a *cis* window of 1 Mb on either side of the transcription start site (TSS) of autosomal and X chromosome genes with a permutation analysis to identify the most significant marker for each gene^42^. We customized the FastQTL software to include an SV for genotype-expression associations when the span of a deletion, duplication, mCNV, or MEI fell within the *cis* window for a particular gene, or when the breakpoints of an inversion or uncharacterized breakend (BND) fell within the *cis* window. For each tissue, we applied a set of covariates including three genotyping principal components, gender, genotyping platform (HiSeq 2000 or HiSeq X Ten), and a variable number of PEER (probabilistic estimation of expression residuals) factors determined by number of samples per tissue, N (N < 150: 15 PEERs, 150 ≤ N ≤ 50: 30 PEERs, N ≥ 250: 35 PEERs)^43^. Note that PEER factor sample sizes include RNA-seq data from individuals lacking WGS, providing more samples for PEER correction than the 147 individuals in the remainder of this study. We performed gene level multiple-testing correction for each of the SV-only and joint analyses using Benjamini-Hochberg at a 10% false discovery rate (FDR).

### Fine mapping of causal variants at eQTLs

We used CAVIAR to untangle linkage disequilibrium to predict a causal variant for each eQTL^18^. CAVIAR assesses summary statistics in conjunction with LD across an associated locus to rank the causal probability of each variant in a region. Thus, we ran FastQTL once again on the 24,780 eQTLs that had previously met FDR thresholds in either the SV-only or joint cis-eQTL mapping analyses to generate the nominal t-statistic for every common variant in the *cis* window. For each eQTL, we selected the most significant SV as well as the 100 most significant SNVs or indels (based on nominal p-value) in the *cis* window and estimated their pairwise LD using linear regression. For SVs, we used allele balance rather than discrete genotype for computing LD. We ran CAVIAR at each of these eQTLs using the t-statistics and signed R values of LD among the 101 variants with a causal set size of 1.

As an alternate estimate of the causal role of structural variation, at each eQTL discovered by either the SV-only or joint analyses, we applied a linear mixed model (LMM) to partition the heritability of each eGene’s expression into a fixed effect from the SV, and a random effect representing the cumulative heritability of the 1,000 most significant SNPs and indels in the *cis* region. This method mirrors that of several other studies that have examined relative contributions of distinct variant classes on a quantitative trait in n individuals^19–21^. We first corrected for the same covariates as in the cis-eQTL mapping analyses above by linear regression residualization, and then applied a linear model of the form

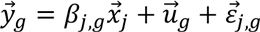

where 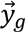 is a vector of the normalized expression values, *β_j,g_* is the effect of allele dosage of SV j on gene g, *x_j_* is a n-length vector of genotypes at SV j, 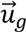 is a n-length vector of random effects drawn from the genetic relatedness matrix (GRM) with 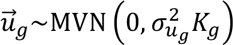, and 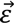 is a random error term drawn from 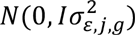 representing unexplained variance. We defined the n x n dimensional GRM (K_g_) with entries 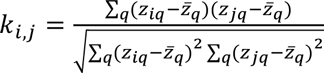for the 1,000 SNV and indel variants (*z_q_*) that are most significantly associated with the expression of eGene *g*.

Solving this equation with GCTA produces an estimate of variance where (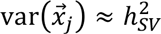).

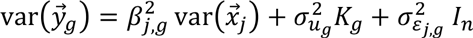

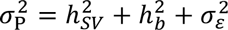

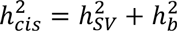

Heritability estimates for a small number of eQTLs could not be calculated due to non-positive definite matrices likely arising from small sample sizes (808/23,441 of joint eQTLs and 930/24780 of eQTLs detected by either SV-only or joint mapping). These loci were excluded from the heritability analysis and composite causality scores described below. To estimate the fraction of *cis* heritability attributable to SVs across all eQTLs in our data set, we counted the number of eQTLs where 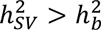 as a fraction of joint eQTLs at which the overall heritability (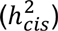) was at least 0.05.

We combined the CAVIAR and heritability estimates of causality into a single composite score for each eQTL by taking the product of the CAVIAR causal probability and the fraction of heritability attributed to the 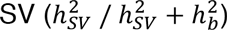. To bound the heritability fraction between 0 and 1, we set the minimum 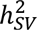 and 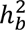 to 10^−6^ and the maximum to 1 before taking the quotient. For SVs that were associated with multiple eQTLs, including those that were independently ascertained in multiple tissues, we selected the eQTL (tissue, gene pair) in which the SV had the highest causality score, resulting in a set of 4,485 distinct SVs.

Though we calculated causality scores (CAVIAR, heritability, composite) for all eQTLs detected by the SV-only or joint analyses, we restricted estimates comparing the relative contributions and effect sizes of SVs, SNVs, and indels to only those 23,441 detected by the joint analysis to eliminate confounding differences in statistical power between the eQTL mapping runs.

### Feature enrichment

We performed intersections between SVs across the range of composite causality score quantiles and various annotated genomic features (**Fig. 3, Supplementary Fig. 1**). We first allocated SVs into 7 bins by the quantile of their composite causality score and then counted the number that intersected with each feature, allowing 1 kb of flanking distance except for the following: exon-altering plot, no flanking distance; proximity to TSS, 10 kb of directional flanking distance; GENCODE genes, no flanking distance; GENCODE exons, no flanking distance; and topologically associated domain boundaries, 5 kb of flanking distance. SVs involved in multiple eQTLs were considered to touch an eGene if they overlapped the exons of genes at any of those eQTLs. SVs from each causality bin were shuffled with BEDTools into non-gapped regions of the genome within 1 Mb of a gene transcription start site^44^. We calculated the fold enrichment of observed feature intersections compared to the median of 100 random shuffled sets of the elements of each bin to control for each bin’s composition of variant types and size distributions. The 95% confidence intervals were derived from the empirical distributions of feature intersections from the of the shuffled set for each bin.

Enhancer positions were defined as those in the Dragon ENhancers database (DENdb) with a minimum support of 2^24^ (Fig. 3b). Positions 10 kb upstream and downstream of the TSS were defined from GENCODE v19 gene positions (Fig. 3c,d), and FunSeq 2.1.0 regions were downloaded from the author’s website (http://archive.gersteinlab.org/funseq2.1.0_data) ^23^. Topologically associated domain boundaries from human embryonic stem cell lines were downloaded from http://compbio.med.harvard.edu/modencode/webpage/hic/hESC_domains_hg19.bed^45^. All other regions were defined by the ENCODE project and downloaded from the UCSC genome browser^22,46^.

### eQTL linkage to GWAS hits

We defined a set of phenotype-associated SNPs from the GWAS catalog v1.0.1 (downloaded 2016-02-04) ^26^. We selected for results with a p-value better than 5 × 10^−8^ and tested in Europeans. When multiple markers within a 100 kb window met this criteria in a single study and a single phenotype, we selected the most significant marker in the window to reduce redundancy, resulting in a set of 4,951 SNPs, of which 4,874 were genotyped in our cohort of 147 samples. We calculated LD between these GWAS hits and variants in our cohort using a linear regression to approximate R^2^, using allele balance rather than discrete genotypes for SVs detected with LUMPY.

### Rare variant association with expression outliers

We began by defining gene expression outliers in each of 544 individuals with RNA-seq data across the 44 tissues available from the GTEx project. Since quantile normalization of expression values (as applied in *cis*-eQTL mapping) can reduce the signal from true expression outliers, we derived PEER-corrected expression values without quantile normalization to define expression outliers. For each tissue, we filtered for genes on the autosomes or the X chromosome in which at least 10 individuals had an RPKM (reads per kilobase of transcript per million mapped reads) > 0.1 and raw read counts > 6. Next we took the log_2_(RPKM + 2) transformation of the data, followed by Z-transformation across each gene. We then removed PEER factors by linear regression residualization (using the same number of factors per tissue as described above, see *Common eQTL mapping)* followed by a final Z-transformation.

We then subsampled the 544 individuals above to select the 117 who were of Caucasian ancestry (since this was the largest subpopulation in our cohort) and had available WGS sequence data. Among these 117 individuals, we identified (sample, gene) pairs at which an individual’s absolute median Z-score of a gene’s expression was at least 2, and there were at least 5 tissues with available expression data for the gene. This amounted to 5,047 gene expression outliers (median: 30 per person, range: 10-298). Next, we identified rare variants that were present in at most two individuals in our cohort of 147 individuals and positively genotyped in at least one of the 117 Caucasian individuals, amounting to 5,660,256 rare variants (4,691 SVs, 4,830,727 SNVs, and 824,836 indels).

We counted the number of rare SVs, SNVs, and indels that co-occurred in the outlier individual that resided within the outlier transcript or 5 kb of flanking sequence. To define the frequency that this occurs by chance, we performed 1,000 random permutations of the outlier individual names in our set to determine the number of rare variants of each type that co-occur with an outlier in a random individual. Notably, we retained the relative number of outliers per individual in each permutation so that individuals with many outliers were still over-represented in the permuted sets.

We performed two reciprocal measures of enrichments. The “outlier-centric” approach tested for a significant difference in the number of outliers that had a rare variant within 5 kb (**Fig. 5a, red points**). However, for SVs in particular, a single large variant may be in proximity to many outliers, and we controlled for this phenomenon with the “variant-centric” approach to test for a significant difference in the number of rare variants that had an outlier gene within 5 kb (**Fig. 5a, blue points**).

To judge the enrichment across thresholds of variant functional impact, we computed a predicted impact score with CADD v1.2 for all variants in our data set^37^. For SVs, we used the highest-scoring base across the affected interval and the confidence intervals around the SV breakpoints. We then computed to the percentile of these impact scores for SVs, SNVs, and indels separately across the full set of allele frequencies. We show the fold-enrichment for the outlier-centric (Supplementary Fig. 7a-c) and variant-centric (Supplementary Fig. 5d-f) approaches across the range of impact score percentiles for each variant class.
To estimate the relative contribution of each variant type to expression outliers, we first defined the fraction of outliers with a likely genetic basis. Across 1,000 shuffled permutations of the data, we observed a median of 1,974 outliers (95% CI: 1,914-2,036) with a rare variant of any type in the outlier individual. We identified 2,417 outliers with a rare variant, representing a net excess of 443 over the expected value (95% CI: 381-503). Thus, of the 5,047 total outliers in our data set, an estimated 8.8% (95% CI: 7.5-10.0%) of outliers have a genetic basis.

We then apportioned these 8.8% of genetically determined outliers amongst SVs, SNVs, and indels according to the net excess of observed outliers within 5 kb of each variant types (**Supplementary Fig. 8**). For outliers that were within 5 kb of multiple variant types (overlaps on the Venn diagram), we allocated the net excess percentage based on the relative strength of their overall fold-enrichment. To achieve this, we first estimated the fraction of expression outliers with a genetic basis within each variant type *(T = {SV,SNV, indel})* as 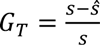, where *s* is the number of observed outliers with 5 kb of a rare variant of type *T* and *ŝ* is the median from the permuted sets (*G_sv_* = 0.94; *G_SNV_* = 0.12; *G_indel_* 0.19 in our data set)‥ Then, for each overlapping region of the Venn diagram, we multiplied the net excess by 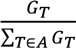 for each of the variant types in each Venn diagram area *A*.

To identify complex variants, we clustered high-confidence rare SVs with breakpoint evidence located no more than 100 kb away from each other and present in the same individual(s). Rare SVs with only read-depth support were not included in this clustering because of their imprecise boundaries. We joined separate clusters if they contained two sides of the same uncharacterized BND. Clusters containing SVs that were previously found to be associated with outlier gene expression were reclassified as a complex deletion, complex duplication, or balanced complex rearrangement by manual curation. During this manual curation, rare SVs with only read-depth support and present in the appropriate sample(s) were added to the rare variant cluster if they overlapped other variants included in the cluster. Upon manual inspection, one outlier-associated SV (LUMPY_BND_195398) with inverted breakpoint orientation was visually determined to have amplified read-depth over the interval and thus reclassified as a complex duplication.

## References

1. Edwards, S. L., Beesley, J., French, J. D. & Dunning, A. M. Beyond GWASs: Illuminating the Dark Road from Association to Function. The American Journal of Human Genetics 93, 779–797 (2013).

2. Lappalainen, T. et al. Transcriptome and genome sequencing uncovers functional variation in humans. Nature 501, 506–511 (2013).

3. The GTEx Consortium et al. The Genotype-Tissue Expression (GTEx) pilot analysis: Multitissue gene regulation in humans. Science 348, 648–660 (2015).

4. Battle, A. et al. Characterizing the genetic basis of transcriptome diversity through RNA-sequencing of 922 individuals. Genome Research 24, 14–24 (2014).

5. 1000 Genomes Project Consortium et al. A global reference for human genetic variation. Nature 526, 68–74 (2015).

6. Conrad, D.F. et al. Origins and functional impact of copy number variation in the human genome. Nature 464, 704–712 (2010).

7. Alkan, C., Coe, B.P. & Eichler, E.E. Genome structural variation discovery and genotyping. Nat. Rev. Genet. 12, 363–375 (2011).

8. Weischenfeldt, J., Symmons, O., Spitz, F. & Korbel, J.O. Phenotypic impact of genomic structural variation: insights from and for human disease. Nat. Rev. Genet. 14, 125–138 (2013).

9. Stranger, B.E. et al. Relative impact of nucleotide and copy number variation on gene expression phenotypes. Science 315, 848–853 (2007).

10. Schlattl, A., Anders, S., Waszak, S.M., Huber, W. & Korbel, J.O. Relating CNVs to transcriptome data at fine resolution: assessment of the effect of variant size, type, and overlap with functional regions. Genome Research 21, 2004–2013 (2011).

11. Sudmant, P.H. et al. An integrated map of structural variation in 2,504 human genomes. Nature 526, 75–81 (2015).

12. Layer, R.M., Chiang, C., Quinlan, A.R. & Hall, I.M. LUMPY: A probabilistic framework for structural variant discovery. Genome B/ol 15, R84 (2014).

13. Chiang, C. et al. SpeedSeq: ultra-fast personal genome analysis and interpretation. Nat Meth 12, 966–968 (2015).

14. Handsaker, R.E., Korn, J.M., Nemesh, J. & McCarroll, S.A. Discovery and genotyping of genome structural polymorphism by sequencing on a population scale. Nature Genetics 43, 269–276 (2011).

15. Danecek, P. et al. The variant call format and VCFtools. Bioinformatics 27, 2156–2158 (2011).

16. Ongen, H., Buil, A., Brown, A.A., Dermitzakis, E.T. & Delaneau, O. Fast and efficient QTL mapper for thousands of molecular phenotypes. Bioinformatics 32, 1479–1485 (2016).

17. McKenna, A. et al. The Genome Analysis Toolkit: A MapReduce framework for analyzing next-generation DNA sequencing data. Genome Research 20, 1297–1303 (2010).

18. Hormozdiari, F., Kostem, E., Kang, E.Y., Pasaniuc, B. & Eskin, E. Identifying causal variants at loci with multiple signals of association. Genetics 198, 497–508 (2014).

19. Gymrek, M. et al. Abundant contribution of short tandem repeats to gene expression variation in humans. Nature Genetics 48, 22–29 (2016).

20. Yang, J., Lee, S.H., Goddard, M.E. & Visscher, P.M. GCTA: a tool for genome-wide complex trait analysis. Am. J. Hum. Genet. 88, 76–82 (2011).

21. Gusev, A. et al. Partitioning heritability of regulatory and cell-type-specific variants across 11 common diseases. Am. J. Hum. Genet. 95, 535–552 (2014).

22. Consortium, T. E. P. et al. An integrated encyclopedia of DNA elements in the human genome. Nature 488, 57–74 (2012).

23. Fu, Y. et al. FunSeq2: a framework for prioritizing noncoding regulatory variants in cancer. Genome Biol 15, 153 (2014).

24. Ashoor, H., Kleftogiannis, D., Radovanovic, A. & Bajic, V.B. DENdb: database of integrated human enhancers. Database 2015, (2015).

25. Nicolae, D.L. et al. Trait-Associated SNPs Are More Likely to Be eQTLs: Annotation to Enhance Discovery from GWAS. PLoS Genet6, e1000888 (2010).

26. Welter, D. et al. The NHGRI GWAS Catalog, a curated resource of SNP-trait associations. Nucleic Acids Research 42, D1001–6 (2014).

27. Gretarsdottir, S. et al. Genome-wide association study identifies a sequence variant within the DAB2IP gene conferring susceptibility to abdominal aortic aneurysm. Nature Genetics 42, 692–697 (2010).

28. Okada, Y. et al. Genetics of rheumatoid arthritis contributes to biology and drug discovery. Nature 506, 376–381 (2013).

29. Suzuki, A. et al. Functional haplotypes of PADI4, encoding citrullinating enzyme peptidylarginine deiminase 4, are associated with rheumatoid arthritis. Nature Genetics 34, 395–402 (2003).

30. Yang, X.-K. et al. Associations Between PADI4 Gene Polymorphisms and Rheumatoid Arthritis: An Updated Meta-analysis. Archives of Medical Research 46, 317–325 (2015).

31. Wu, C. et al. Joint analysis of three genome-wide association studies of esophageal squamous cell carcinoma in Chinese populations. Nature Genetics 46, 1001–1006 (2014).

32. Barrett, J.H. et al. Genome-wide association study identifies three new melanoma susceptibility loci. Nature Genetics 43, 1108–1113 (2011).

33. de Cid, R. et al. Deletion of the late cornified envelope LCE3B and LCE3C genes as a susceptibility factor for psoriasis. Nature Genetics 41, 211–215 (2009).

34. Chambers, J.C. et al. Genome-wide association study identifies loci influencing concentrations of liver enzymes in plasma. Nature Genetics 43, 1131–1138 (2011).

35. Li, X. & Montgomery, S.B. Detection and Impact of Rare Regulatory Variants in Human Disease. Front. Genet. 4, (2013).

36. Li, X. et al. Transcriptome sequencing of a large human family identifies the impact of rare noncoding variants. Am. J. Hum. Genet. 95, 245–256 (2014).

37. Kircher, M. et al. A general framework for estimating the relative pathogenicity of human genetic variants. Nature Publishing Group 46, 310–315 (2014).

38. Li, H. Aligning sequence reads, clone sequences and assembly contigs with BWA-MEM. arXiv.org 1303.3997 (2013).

39. Trapnell, C., Pachter, L. & Salzberg, S.L. TopHat: discovering splice junctions with RNA-Seq. Bioinformatics 25, 1105–1111 (2009).

40. Harrow, J. et al. GENCODE: producing a reference annotation for ENCODE. Genome Biol 7, S4–9 (2006).

41. Deluca, D.S. et al. RNA-SeQC: RNA-seq metrics for quality control and process optimization. Bioinformatics 28, 1530–1532 (2012).

42. Ongen, H., Buil, A., Brown, A., Dermitzakis, E. & Delaneau, O. Fast and efficient QTL mapper for thousands of molecular phenotypes. bioRxiv (2015). doi:10.1101/022301

43. Stegle, O., Parts, L., Piipari, M., Winn, J. & Durbin, R. Using probabilistic estimation of expression residuals (PEER) to obtain increased power and interpretability of gene expression analyses. Nat Protoc 7, 500–507 (2012).

44. Quinlan, A.R. & Hall, I.M. BEDTools: a flexible suite of utilities for comparing genomic features. Bioinformatics 26, 841–842 (2010).

45. Ho, J. et al. Comparative analysis of metazoan chromatin organization. Nature (2014).

46. Kent, W.J. et al. The Human Genome Browser at UCSC. Genome Research 12, 996–1006 (2002).

